# Interpretable Solutions for Stochastic Dynamic Programming

**DOI:** 10.1101/2024.08.05.606713

**Authors:** Jonathan Ferrer-Mestres, Thomas G. Dietterich, Olivier Buffet, Iadine Chadès

## Abstract

1. In conservation of biodiversity, natural resource management and behavioural ecology, stochastic dynamic programming, and its mathematical framework, Markov decision processes (MDPs), are used to inform sequential decision-making under uncertainty. Models and solutions of Markov decision problems should be interpretable to derive useful guidance for managers and applied ecologists. However, MDP solutions that have thousands of states are often difficult to understand. Difficult to interpret solutions are unlikely to be applied, and thus we are missing an opportunity to improve decision-making. One way of increasing interpretability is to decrease the number of states.
2. Building on recent artificial intelligence advances, we introduce a novel approach to compute more compact representations of MDP models and solutions as an attempt at improving interpretability. This approach reduces the size of the number of states to a maximum number *K* while minimising the loss of performance compared to the original larger number of states. The reduced MDP is called a *K*-MDP. We present an algorithm to compute *K*-MDPs and assess its performance on three case studies of increasing complexity from the literature. We provide the code as a MATLAB package along with a set of illustrative problems.
3. We found that *K*-MDPs can achieve a substantial reduction of the number of states with a small loss of performance for all case studies. For example, for a conservation problem involving Northern Abalone and Sea Otters, we reduce the number of states from 819 to 5 states while incurring a loss of performance of only 1%. For a dynamic reserve selection problem with seven dimensions, while an impressive reduction in the number of states was achieved, interpreting the optimal solutions remained challenging.
4. Modelling problems as Markov decision processes requires experience. While several models may represent the same problem, reducing the number of states is likely to make solutions and models more interpretable and facilitate the extraction of meaningful recommendations. We hope that this approach will contribute to the uptake of stochastic dynamic programming applications and stimulate further research to increase interpretability of stochastic dynamic programming solutions.

## 1 Introduction

Stochastic dynamic programming (SDP) is a powerful solution technique to find the best time-dependent decisions in the presence of uncertainty about the dynamics of a system (Bellman, 1957). SDP is applied across a broad range of decision-making applications from behavioural ecology and natural resource management to conservation of biodiversity and epidemiology (McCarthy et al., 2001; Marescot et al., 2013; Boettiger et al., 2015; Schwarz et al., 2016; Westphal et al., 2003; Milner-Gulland, 1997). While SDP refers to the optimisation technique, Markov Decisions Processes (MDPs) is the mathematical formalism employed to model many sequential decision-making problems(Bellman, 1957; Puterman, 2014). MDPs can model problems where the dynamics of the system are stochastic and the state of the system is completely observable (Marescot et al., 2013). In MDPs, a state is a collection of variables with a given value, for example, the abundance of a species at a site. Actions represent the decisions that can be implemented at each time step, and transition functions represent the probability of transitioning from one state to another state when an action is implemented. Finally, a reward function specifies how to evaluate states and actions. Some states might be more desirable than others. For example high abundance states are more valuable in conservation of threatened species than low abundance states, while actions often come at a cost. In MDP problems, the objective is usually to find the state-dependent actions that will maximise the expected sum of rewards over time (i.e., maximise the chance of the system being in high reward states).

In ecology, MDP models have been developed for many problems including population recovery of threatened species under limited resources (Chades et al., 2012), re-introducing captive-raised species (Collazo et al., 2013), controlling invasive species (Moore et al., 2010), reducing the risk of disease spread (Péron et al., 2017), optimally allocating conservation resources between regions (Wilson et al., 2006), managing fisheries (Memarzadeh et al., 2019), adaptively managing natural resources (Pozzi et al., 2017), managing habitats (Johnson et al., 2011) and testing behavioral ecology theories (Venner et al., 2006; Mangel et al., 1988) to cite a few. One of the barriers identified as preventing a greater uptake of MDPs is the complexity of understanding their models and optimised solutions (Tulloch et al., 2015). In human-operated systems, where the domain expert decides whether a recommendation proposed by a system is appropriate or not, it is critical that models and solutions can be interpreted and explained (Petrik and Luss, 2016). Otherwise, for most non-simple decision problems, users tend to apply simple decision-making strategies (heuristics) instead of attempting to find and understand an optimal solution (Reeson and Dunstall, 2009).

Traditionally, research in artificial intelligence (AI) has focused on developing algorithms (Sigaud and Buffet, 2013) to efficiently solve MDP problems for autonomous agents, such as self-driving cars (Brechtel et al., 2011) and robots (Kober et al., 2013) rather than increasing interpretability. With the growing number of MDP applications to human-operated domains, such as conservation of biodiversity and natural resources management, developing approaches that provide easy-to-interpret solutions is critical to increase uptake of MDPs as a modelling framework and to trust solutions computed by MDP algorithms. A possible path forward is to reduce the number of states to produce more compact solutions that are easier to understand and interpret by human operators (Nicol and Chadès, 2012). Reducing the number of states in MDPs has been explored before, early in the 90’s, but with the aim of solving problems with large state spaces rather than increasing interpretability (see (Dean et al., 1997; Li et al., 2006; Abel et al., 2016, 2018) and supp. info.). However, those approaches are not human-oriented, and they do not give managers control over how much the state space should be reduced. To address these shortcomings, we build on a recent AI model, *K*-MDP, and algorithms that allow a user to trade-off performance and the maximum number of states *K* in the *K*-MDP (Ferrer-Mestres et al., 2020).

The principle of these algorithms consists in reducing the number of states of an MDP problem by grouping states into bins while minimising the loss of performance between the original MDP solution and the reduced optimal *K*-MDP solution, where *K* represents the maximum number of states. To gain insights into the usefulness of these algorithm in ecology, we assess *K*-MDPs on three case studies of increasing complexity: managing a wolf population with one state variable (Marescot et al., 2013), recovering two endangered species, sea otter and northern abalone, with two state variables (Chades et al., 2012), and a dynamic reserve site selection problem with seven state variables (Costello and Polasky, 2004; Sabbadin et al., 2007; Chadès et al., 2014). We evaluate the interpretability of the resulting models and solutions, and provide recommendations for future research. While this approach has a lot of potential to increase interpretability of MDP solutions, more needs to be done.

## 2 Materials and Methods

### 2.1 Markov Decision Processes

For ease of presentation of MDPs, we will assume a hypothetical conservation problem where the objective is to maximise the abundance of a threatened species over time. The species is threatened by an invasive predator and by illegal harvesting. Every six months, managers survey the population and have close-to-perfect assessment of the population abundance of the threatened species (state variable). Managers can decide either implementing anti-poaching measures or culling the population of the invasive predator for the next six months (two actions). These actions come at the same cost and have probabilistic outcomes. To help managers best invest their limited resources, we can model this problem as an MDP and provide recommendations based on the current abundance of the threatened species. MDPs have three key characteristics:

1. The current state of the system is completely observable: we can perfectly measure the abundance of the current populations.
2. The dynamics of the system can be stochastic: an action has probabilistic outcomes.
3. The dynamics of the system is assumed Markovian: the probability of transitioning to the next state at time *t* + 1, only depends on the state and action implemented at time *t*.

Formally, MDPs are a mathematical framework for modeling sequential decision-making problems under uncertainty (Bellman, 1957; Puterman, 2014). An MDP is defined as a tuple ⟨*S, A, T, r, γ*⟩, where:

- *S* is a finite set of states. Here, *s* ∈ *S* represents the abundance of the threatened species;
- *A* is a finite set of actions from which the manager must choose an action *a* at each time step; e.g., *A* ={anti-poaching, culling};
- *T* is the Markovian transition probability function describing the stochastic dynamics of the system; an element *T* (*s, a, s*^*′*^) = *p*(*s*′|*s, a*) represents the probability of being in state *s*′ at time *t* + 1 if action *a* has been applied in state *s* at time *t*; e.g., the probability of increasing the abundance of species after an anti-poaching measure has been implemented;
- *r* is the reward function describing the immediate benefits and costs of applying an action in a given state; an element *r*(*s, a*) represents the reward or cost of applying action *a* in state *s* at time *t*; e.g., the cost of applying culling measures;
- *γ* is a discount factor; it can take a value between 0 and 1; a discount factor less than 1 means that immediate rewards are more valuable than later ones (Koopmans, 1960).

A solution to an MDP is a function *π* : *S* → *A*, called a policy, that maps states into actions. A policy specifies what action to take in each state. From a given state *s*, the decision-maker applies an action *a*, receives a reward *r*(*s, a*), and the state of the system is updated to *s*^*′*^ with probability *p*(*s*^*′*^|*s, a*).

#### Policy representation

Of interest to increasing interpretability of MDP solutions is the visualisation of a policy. A popular way of visualizing policies is the policy graph, a visual representation where nodes represent states and edges represent the optimal action to apply and the probability of transitioning to another state. Figure 1 shows a small part of a policy graph involving one state *s*, one action *a* = *π*(*s*), and two possible transitions from state *s* to state *s*^*′*^ or *s*^*′′*^ with corresponding probabilities *p*(*s*^*′*^|*π*(*s*), *s*) and *p*(*s*^*′′*^|*π*(*s*), *s*) respectively. Figure 2 shows an example policy graph for a conservation decision problem with two endangered species, the sea otter and the northern abalone, involving 819 states (Chades et al., 2012). This graph is obviously impossible to interpret because of the number of states (819). The aim in this paper is to increase the interpretability of MDP models and solutions.

**Figure 1:**
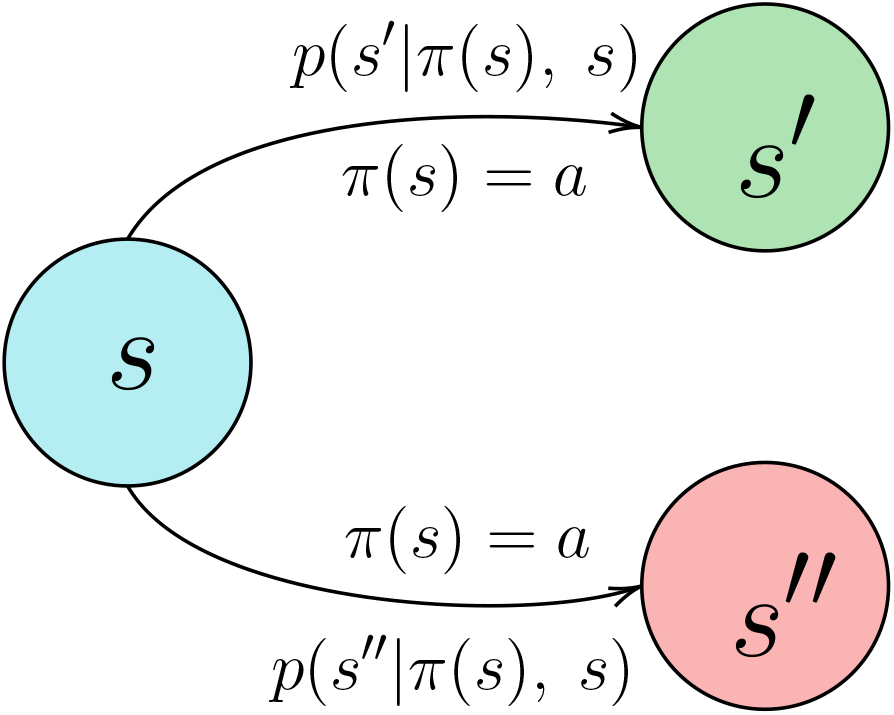
Policy graph where nodes represent states, and arrows represent possible transitions to next states. *s* is the current state; *π*(*s*) = *a* is the action to apply in state *s*, following policy *π*; *s*^*′*^ and *s*^*′′*^ are the resulting states of applying action *π*(*s*) in *s*; *p*(*s*^*′*^|*π*(*s*), *s*) and *p*(*s*^*′′*^|*π*(*s*), *s*) are the transition probabilities.

**Figure 2:**
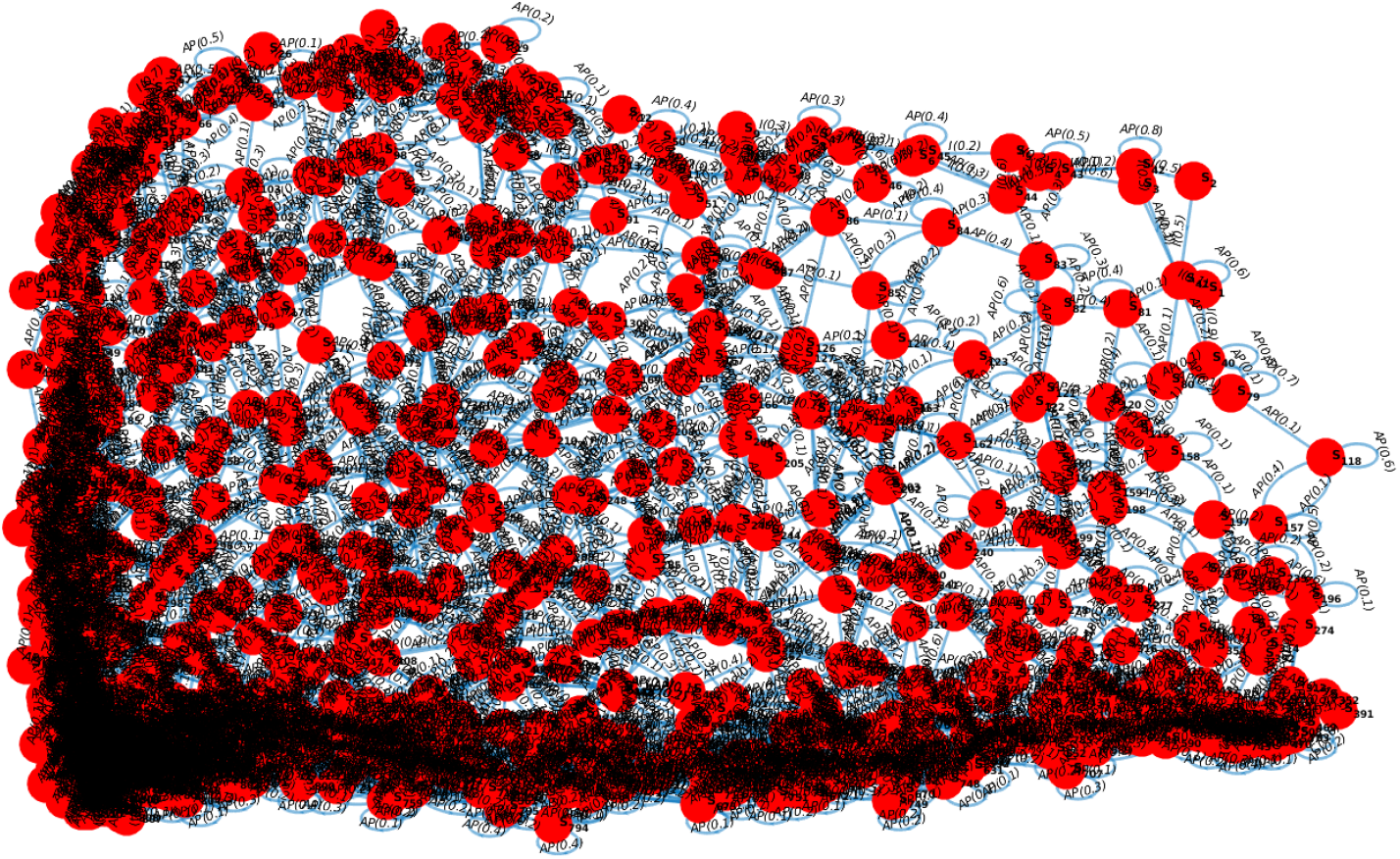
Policy graph representing an optimal strategy for a conservation problem of two endangered species(Chades et al., 2012). Nodes represent the 819 states and edges represent the optimal action to take and its probability to transition to another state (up to 819^2^ edges).

#### Optimisation criterion

The first step when defining a decision problem is to formulate an optimisation criterion. In this paper, the optimisation criterion is to maximise the expected sum of discounted rewards over an infinite time horizon: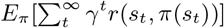, where *s*_*t*_ denotes the state and *π*(*s*_*t*_) = *a*_*t*_ represents the action implemented at time *t* when implementing policy *π*.

#### Value function

The value of a policy *π* in state *s* is denoted *V* ^*π*^(s), and it represents the expected sum of discounted rewards of implementing policy *π* starting in state *s* and continuing to a time horizon *H* = ∞. We can write this as

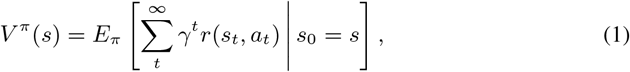

for every state *s* ∈ *S* (Bellman, 1957). There exists at least one optimal policy *π*^∗^ that maximises *V* ^*π*^(*s*) over all states *s* ∈ *S*. Let *V* ^∗^ denote the optimal value function of an optimal policy *π*^∗^.

Several algorithms have been developed to solve MDPs (Marescot et al., 2013; Fackler, 2011; Chadès et al., 2014), including Value iteration.

#### Value iteration

Value iteration finds the optimal value function *V* ^∗^ by solving Bellman’s dynamic programming equation (Bellman, 1957):

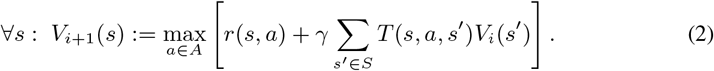

For *i* = 0, the value function *V*_0_ is initialized to an arbitrary value. Then, *V*_*i*+1_ is set to the result of evaluating the right-hand side of Equation 2. As *i* tends to infinity, *V*_*i*_ will converge to the optimal value function *V* ^∗^. In practice, value iteration is halted when the difference between two successive value functions *V*_*i*_ and *V*_*i*+1_ is small (Chadès et al., 2014).

### 2.2 The *K*-MDP approach

Given a number *K, K*-MDP algorithms find a compact version of the original MDP, with at most *K* states, called a *K*-MDP. In this section, we define *K*-MDPs.

#### 2.2.1 Definition

Given an MDP *M* = ⟨*S, A, T, r, γ*⟩, let us define a *K*-MDP (Ferrer-Mestres et al., 2020) as a tuple *M*_*K*_ = ⟨*S*_*K*_, *ϕ, A, T*_*K*_, *r*_*K*_, *γ*⟩, where:

- *S*_*K*_ is the reduced state space of size at most *K*. These states are called *abstract states*;
- *ϕ* is a mapping function from the original MDP state set *S* to the *K*-MDP state set *S*_*K*_, i.e., *ϕ*(*s*) = *s*_*K*_. The inverse function *ϕ*^−1^(*s*_*K*_) maps an abstract state *s*_*K*_ ∈ *S*_*K*_ to its constituent states in the original MDP, i.e., *ϕ*^−1^(*s*_*K*_) = {*s* ∈ *S* | *ϕ*(*s*) = *s*_*K*_ };
- *A* is the same set of actions as in the MDP;
- *T*_*K*_ : *S*_*K*_ × *A* × *S*_*K*_ → [0, 1] is the *K*-MDP probability transition function. An element 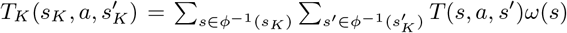, represents the probability of being in state *s*′_*K*_ at time *t* + 1 given action *a* was implemented in state *s*_*K*_ at time *t*. Weights *ω*(*s*) represent a probability distribution over the original states that aggregate to an abstract state 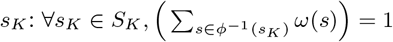;
- *r*_*K*_ : *S*_*K*_ × *A* → [0, *R*_*max*_] is the reward function describing the benefits and costs (dynamical) system; an element 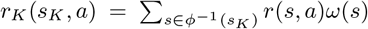 represents the weighted reward or cost of applying action *a* in state *s*_*K*_ at time *t*. In this paper, the weights form a uniform probability distribution over the original states: 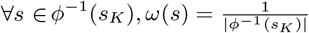;
- *γ* remains unchanged.

An optimal solution for a *K*-MDP is a policy 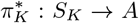 that maximizes the expected sum of discounted rewards and can be calculated using value iteration. Any optimal *K*-MDP policy can be applied to the original MDP by using the mapping function *ϕ*, with the associated value function 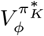, where

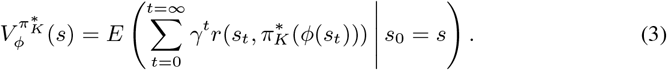

Essentially, 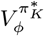 represents the performance of the *K*-MDP policy 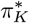 when applied to the original MDP. Many different *K*-MDPs can be constructed from the given MDP, but their performance (measured by their value functions 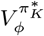) can differ. For example, in Figure 3b, the abstract states can be defined differently by setting 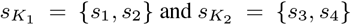. The challenge is to find a *K*-MDP whose optimal policy minimises the loss of performance compared to the optimal policy of the original MDP.

**Figure 3:**
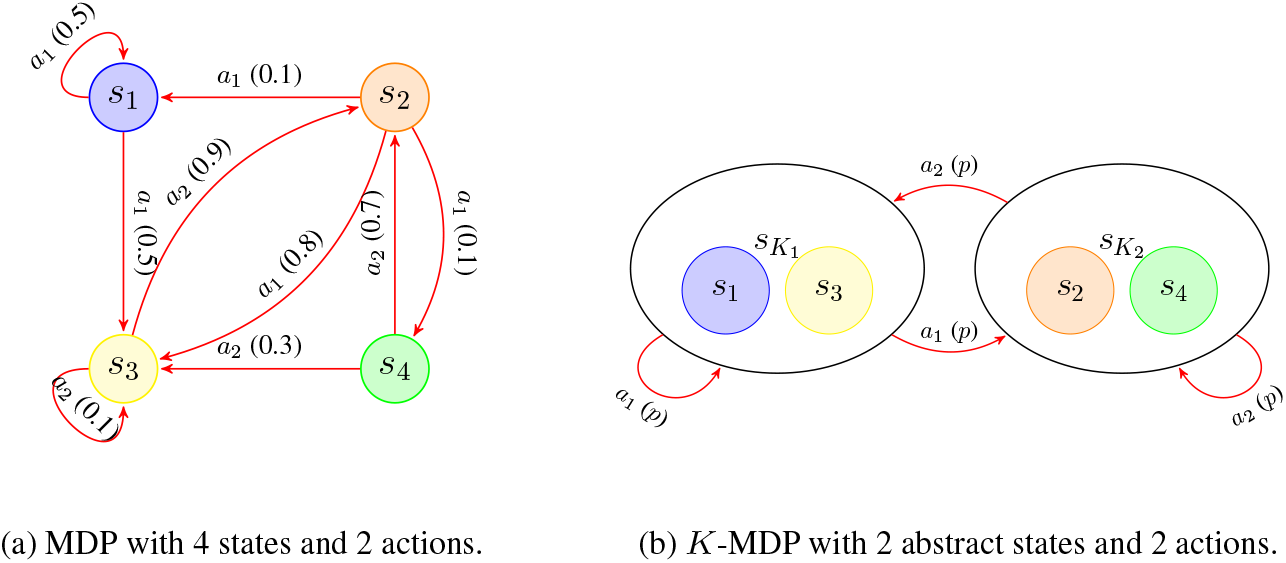
Example of a 4-state MDP and its *K*-MDP with 2 abstract states. *s*_*i*_ are original MDP states; 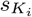 are abstract states. An edge 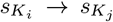 labeled with *a*_*z*_ (*p*) indicates the transition 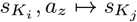 has probability *p*.

For a summary of the different notations used in MDPs and *K*-MDPs, please refer to Supplementary material. Figure 3b is an illustrative example of an MDP and one possible *K*-MDP.

#### 2.2.2 How to calculate a good *K*-MDP?

The problem of finding the best reduced state space *S*_*K*_ is formulated as a gap minimization problem between the original MDP value function and the reduced *K*-MDP value function.

The optimal gap *gap*^∗^ is the difference between the performance of the optimal MDP solution policy *π*^∗^ and the performance of the optimal *K*-MDP policy 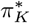, both executed on the original MDP. This difference is measured as the maximum difference (over all states *s*) of the difference in the corresponding value functions:

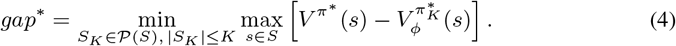

(Ferrer-Mestres et al., 2020) developed an algorithm to compute and solve *K*-MDPs and minimise this gap. To provide an algorithm with performance guarantee to the user, they used a concept developed in artificial intelligence called state abstraction functions (Dean and Givan, 1997; Li et al., 2006). These functions specify how states in the original MDP should be grouped together in the *K*-MDP. This is a challenging design choice, because the quality of the state-grouping rules will influence the performance of the reduced *K*-MDP. Here, we will only present the information required to understand the state-abstraction function of the best *K*-MDP algorithm. Please, refer to the supplementary information or to Ferrer-Mestres et al. (2020) for further details.

#### 2.2.3 State abstraction function

A good state abstraction groups together states that have the same or similar behaviour and minimizes the gap (Equation 4). Ferrer-Mestres et al. (2020) investigated several state abstraction functions; here we will only explain the state abstraction function 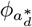 that yielded the most interesting results. This state abstraction function specifies that two states should be grouped together if their optimal actions are the same and they have similar values, i.e., they belong to the same bin or group of values. A state *s* is assigned to a bin by calculating the formula 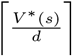, where the notation ⌈*x*⌉, called “ceiling”, denotes the smallest integer that is greater than or equal to *x*; *V* ^∗^(*s*) is the optimal value for the state *s* (Equation 1); and *d >* 0 represents the size of the bin.

Formally, the function 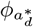 groups two states *s*_1_ and 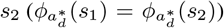, if the following condition holds:

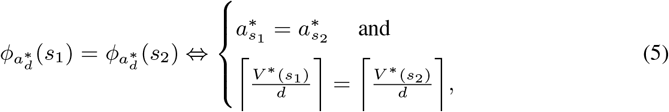

where 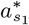 and 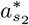 are the optimal actions of states *s*_1_ and *s*_2_. This state abstraction function has valuable mathematical properties that guarantee that the performance loss will be bounded. Interested readers can refer to Ferrer-Mestres et al. (2020).

Table 1 shows an example for an MDP with four states and two actions. In this example, when *d* = 0.15, only states *s*_1_ and *s*_2_ are aggregated, so the resulting *K*-MDP would contain 3 abstract states.

**Table 1:** Illustrative example of ceiling function using 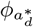 and *d* = 0.15. States *s*_1_, *s*_2_ and *s*_3_ have the same ceil values, however only states *s*_1_ and *s*_2_ can be aggregated because they have the same optimal action.

| State | $V^*(\cdot)$ | $\frac{V^*(\cdot)}{d}$ | $\left\lceil \frac{V^*(\cdot)}{d} \right\rceil$ | $a_s^*$ |
| --- | --- | --- | --- | --- |
| $s_1$ | 0.50 | 3.33 | **4** | **$a_2$** |
| $s_2$ | 0.46 | 3.07 | **4** | **$a_2$** |
| $s_3$ | 0.59 | 3.93 | **4** | $a_1$ |
| $s_4$ | 0.32 | 2.13 | 3 | $a_2$ |

The value of *d* determines the number of bins. If *d* is small, the number of bins will be very large, whereas if *d* is greater than or equal to the maximum possible *V* ^∗^ value, then there will be only one bin per action. The challenge is to choose the value of *d* such that the optimal gap is minimized and the number of abstract states is smaller or equal to *K. A*-*K*-MDP algorithm, finds the best value for *d* using binary search.

#### 2.2.4 Algorithm for solving *K*-MDPs

We now explain the principles of the algorithm (Algorithm 1) introduced by Ferrer-Mestres et al. (2020). This algorithm takes as input an MDP, a state abstraction function 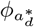, a precision parameter *p*_*target*_, and a constraint on the number of states *K*. The output is a *K*-MDP. The algorithm proceeds in three phases: i) the MDP is solved via value iteration and the optimal policy *π*^∗^ and value function *V* ^∗^ are obtained (Line 1). ii) the abstraction function 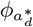 is applied to aggregate the MDP states into *K* abstract states (Line 3-16). iii) the weighting functions *ω*(*s*), abstract transition function *T*_*K*_, abstract reward function *r*_*K*_, abstract optimal policy 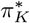 and value function 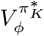 are computed (Line 17).

The abstraction phase applies an aggregation rule that is determined by 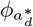 and by a quantization parameter *d*. To find a value of *d* that results in the desired number of states *K*, the *A*-*K*-MDP algorithm performs a binary search. It initializes *d* to the average of the smallest and largest *V* ^∗^ values (Line 2). If this produces too many abstract states, then it increases *d* (Line 12); if it produces too few abstract states, it decreases *d* (Line 14).

The search halts when the optimal value of *d* has been determined to within precision *p*_*target*_ (Line 16). This precision parameter controls the number of iterations when halving *d* (Line 5). Small values of the precision parameter result in small values of *d*. The container *bindings* contains the ceil optimal values and optimal action pairs and, function *group* (Line 10) groups the states that have matching ceil values and optimal actions.

*A*-*K*-MDP can find a feasible *K*-state abstraction because it accounts for the optimal actions in the aggregation process. However, this comes at a cost of not finding a feasible state abstraction when *K* is smaller than the number of actions used in the optimal policy.

Because *A*-*K*-MDP is based on binary search, its computational complexity is attractive, 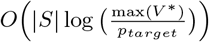. In addition, Ferrer-Mestres et al. (2020) also derived a theoretical bound on the maximum error between the original optimal MDP value function and the reduced optimal *K*-MDP value function (see Table 2).

**Table 2:** Summary of key elements of the *A*-*K*-MDP algorithm: The abstraction function, the aggregation rule, and the theoretical bound on the maximum error (Value Loss) between 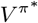 and 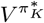.

| $A$-$K$-MDP | |
| --- | --- |
| Function | $\phi_{a_d^*}$ |
| **Aggregation Rule** | $a_{s_1}^* = a_{s_2}^* \quad \text{and} \quad \left\lceil \frac{V^*(s_1)}{d} \right\rceil = \left\lceil \frac{V^*(s_2)}{d} \right\rceil$ |
| **Value Loss** | $\max_{s \in S} V^{\pi^*}(s) - V^{\pi_K^*}(s) \leq \frac{2dR_{max}}{(1-\gamma)^2}$ |

**Table 3:** Summary table of the size reduction and error for each problem and the minimum value of *K* for which we found a suitable state space reduction.

| | $K$ | Size reduction | Error |
| --- | --- | --- | --- |
| Wolf Culling #1<br>$\vert S\vert = 3000, \vert A\vert = 11$ | 6 | 99.83% | 1 % |
| Wolf Culling #2<br>$\vert S\vert = 250, \vert A\vert = 251$ | 50 | 80.00% | 1 % |
| Sea Otter & Northern Abalone<br>$\vert S\vert = 819, \vert A\vert = 4$ | 5 | 99.39% | 1 % |
| Dynamic Reserve site selection<br>$\vert S\vert = 729, \vert A\vert = 6$ | 8 | 98.90% | 0 % |

##### Algorithm 1

*A*-*K*-MDP algorithm

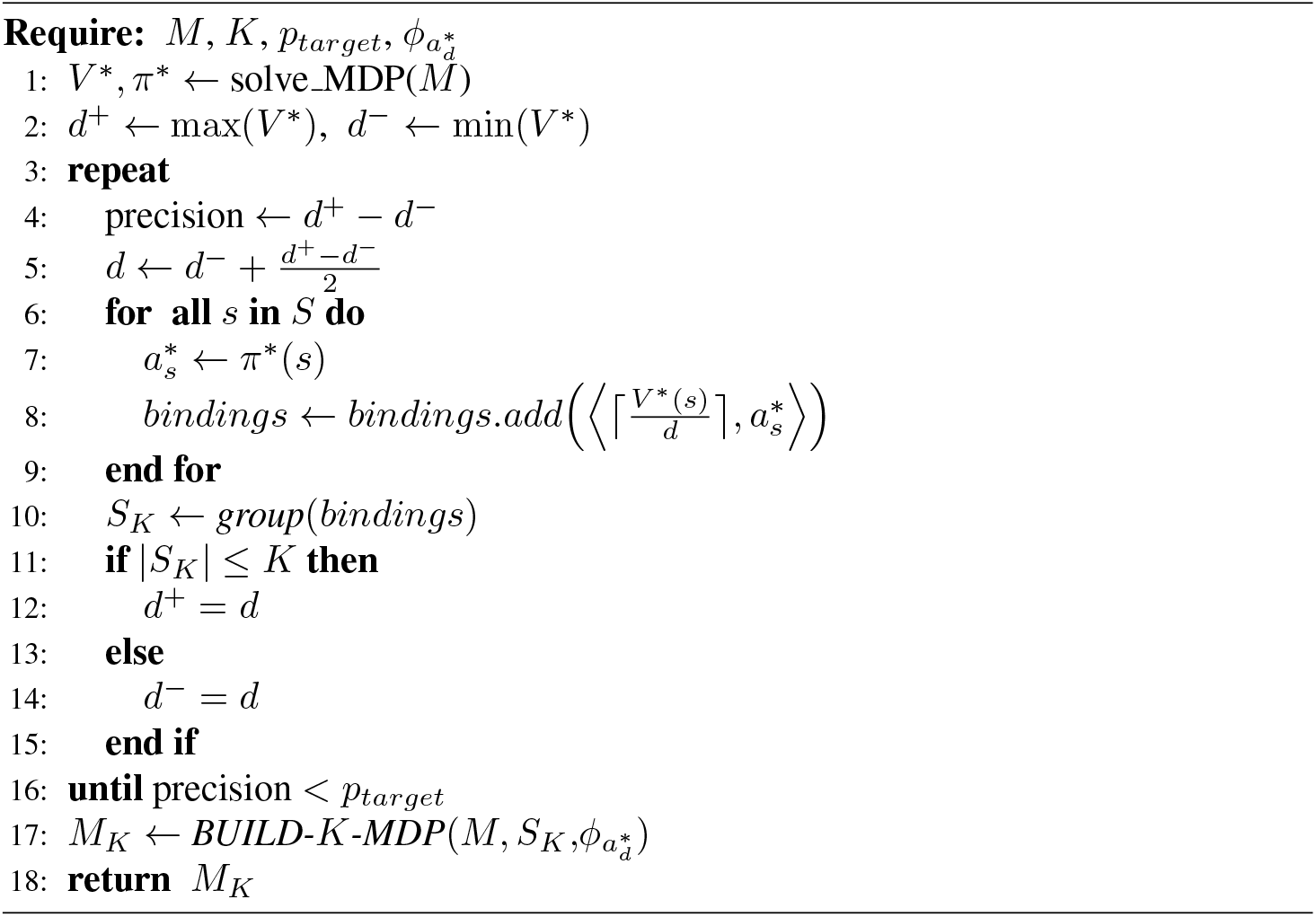

### 2.3 Experimental Evaluation

We assess the performance of *A*-*K*-MDP on three MDP problems from the literature.

#### 2.3.1 Control of a population: Wolf Culling

We first introduce a simple problem with one state variable. The objective of this problem is to maximise the population of the European wolves, ensuring that it remains between *N*_*min*_ and *N*_*max*_ individuals (Marescot et al., 2013). There is only one state variable *X*_*t*_ representing the number of individuals *N*_*t*_ at time *t* and ranging from 0 to *N*_*max*_. There are |*A*| management actions specifying a harvest rate. The harvest rate at time *t* is denoted *H*_*t*_. The dynamics of the system defining the consequences of the actions are derived from an exponential growth population model, assuming an additive effect of human take on total mortality. Rewards increase linearly with abundance when the current state is within [*N*_*min*_, *N*_*max*_].

We propose two instances of this problem. The first instance has 3000 states and 11 actions representing a harvest rate from 0% to 100% with an increment of 10%. States represent the number of individuals. e.g., *s*_1_ represents 0 individuals, *s*_2_ represents 1 individual. The second instance has 250 states, and 251 actions denoting the harvest rate from 0% to 100% with an increment of 1*/*(*N*_*max*_ + 1), allowing as many actions as states. In (Marescot et al., 2013), an optimal policy for this problem was represented graphically as a function of its state variable, the number of individuals (Figure 4).

**Figure 4:**
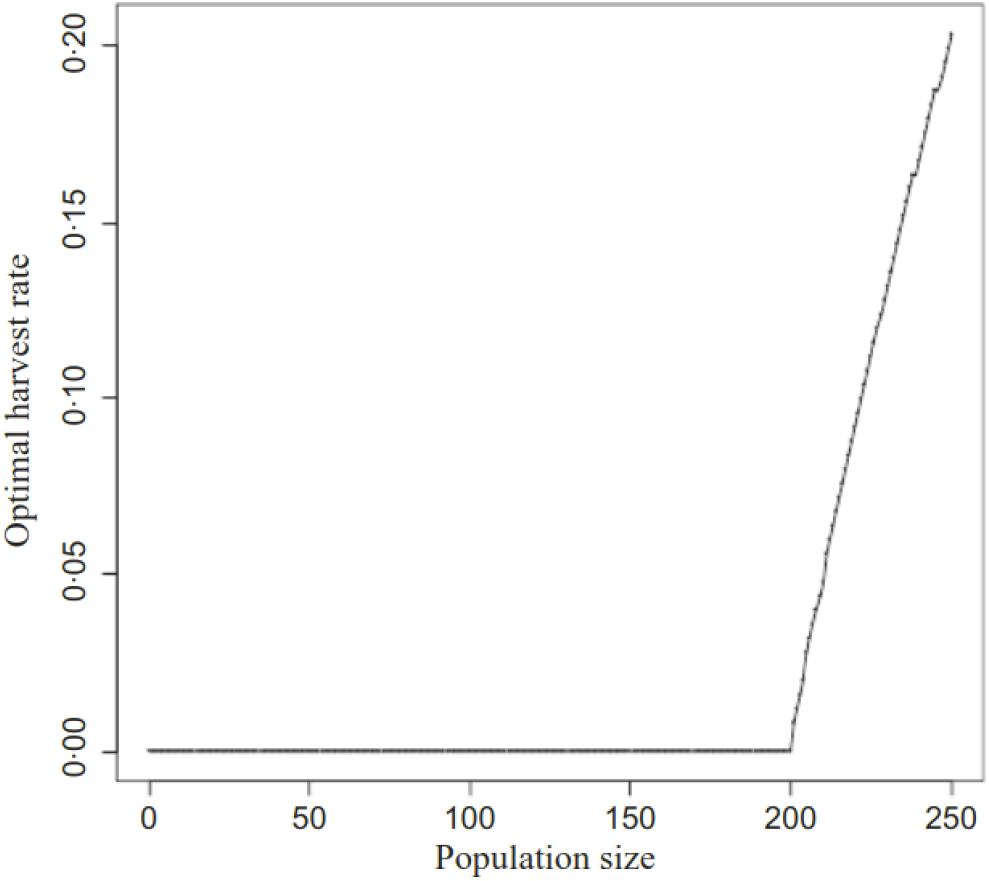
Wolf culling policy represented as a function: *π*(population size) = harvest rate (Marescot et al., 2013).

#### 2.3.2 Recovering two endangered species: Sea Otter and Northern Abalone

Our second problem tackles the recovery of two interacting endangered species (Chades et al., 2012): the sea otter (*Enhydra lutris kenyoni*) and its preferred prey, northern abalone (*Haliotis kamtschatkana*). The authors modeled the interaction between sea otters and northern abalone with three different functional responses: linear, hyperbolic and sigmoid. Here, for simplicity, we focus on a single functional response, the hyperbolic response, which assumes that sea otters mainly feed on abalone. The objective is to maximize the abundance of both species over time. The problem can be modeled as an MDP with 819 states. Each state is composed by 2 state variables: (a) an interval of the density of northern abalone (individuals/*m*^2^) and (b) an interval of the number of sea otters. The initial state *s*_1_ assumes absence of sea otters and an average abalone density of 0.025/*m*^2^. Managers can choose between four different actions at each time step: introduce sea otters, antipoaching measures, control sea otters, and a mix of one half antipoaching measures and one half control sea otters. Sea otters cannot be introduced unless the abalone population has attained a specified recovery goal. The antipoaching action reduces the illegal harvesting of abalone by 50%. The control sea otters action removes the number of otters above 60% of the carrying capacity. The half antipoaching and half control sea otters action takes both actions simultaneously but at half the level of effectiveness. The optimal policies were described in the original manuscript but not represented visually due to their complexities and large sizes (Chades et al., 2012).

Figure 5 shows a simulation of an optimal policy as it was represented by Chades et al. (2012) for the hyperbolic response.

**Figure 5:**
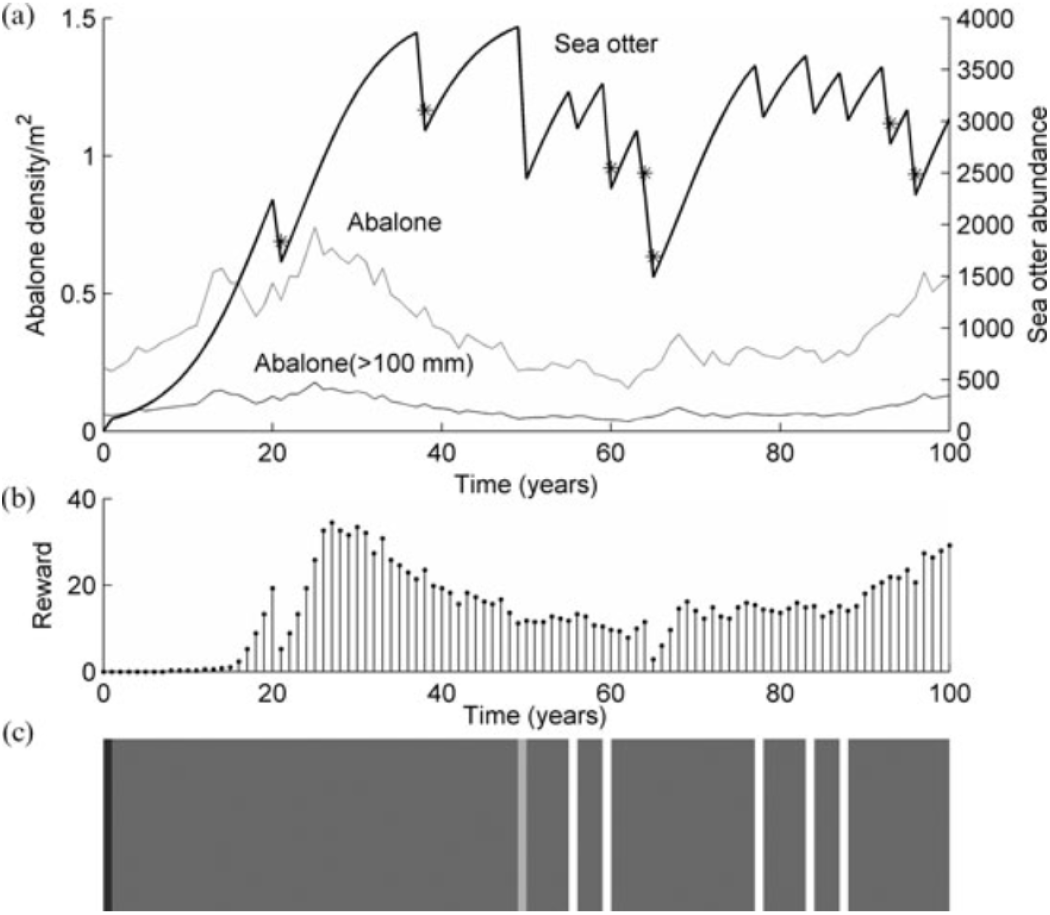
Simulation of the dynamic between sea otters and northern abalone assuming the hyperbolic functional response and applying the optimal policy (a); when both species reach an equivalent level of density or abundance a positive reward is obtained (b); and the optimal policy is: black, introduce sea otters; dark gray, antipoaching; light gray, control of sea otters; white, half antipoaching and half control (c) (Chades et al., 2012).

#### 2.3.3 Dynamic reserve site selection

Our third problem is a dynamic reserve site selection (Chadès et al., 2014). Acquiring land to establish and create reserves is a fundamental action to reduce the loss in biodiversity and protect species. Nonetheless, acquiring all desired sites is virtually always infeasible given the limit on funding (Costello and Polasky, 2004). Managers must decide which available sites should be reserved. This kind of problem has been modeled as an MDP (Costello and Polasky, 2004; Sabbadin et al., 2007). A site can be available, developed or reserved. Every year, the available sites can be developed according to a set probability, and only a limited number of sites can be reserved. The objective is to maximise the number of species conserved over time by reserving available sites to prevent them from being developed.

We employ the model developed by Sabbadin et al. (2007) and coded by Chadès et al. (2014). We assume a matrix *D* of size *J* × *I* representing the distribution of each species *i* = 1, …, *I* at sites *j* = 1, …, *J*. An element *D*_*ji*_ is 1 if site *j* is suitable for species *i* and 0 otherwise. A species *i* exists in site *j* if and only if site *j* is not developed. At any given time *t*, a site *j* can be available (0), reserved (1), or developed (2). There are in total |*S*| = 3^*J*^ states. There are |*A*| = *J* possible actions corresponding to the selection of a site for reservation. A site *j* can only be reserved if it is available. Only one site can be reserved at each time step. Every year, any available and non-reserved site *j* can become developed with probability *p*_*j*_. The objective is to protect the largest number of unique species in reserved sites. For this reason, the reward function is defined as the number of additional unprotected species that become protected when a site is reserved.

For our experiments, we select *J* = 6 sites and *I* = 7 species. The problem has six state variables and the total number of states is |*S*| = 3^*J*^ = 729; there are |*A*| = *J* = 6 actions. States are defined as vectors where each dimension represents a site *j*. In our case, states are 6-dimensional vectors: state = [0, 0, 1, 2, 0, 0] means that sites *j* = 1, 2, 5 and 6 are available, site *j* = 3 is reserved, and site *j* = 4 is developed. At any given time step, an available site can become developed with 0.1 probability. Figure 6 shows a simulation of the optimal policy as represented by Chadès et al. (2014). Representing the entire original optimal policy is far from trivial given the high dimensionality of this problem (six state variables).

**Figure 6:**
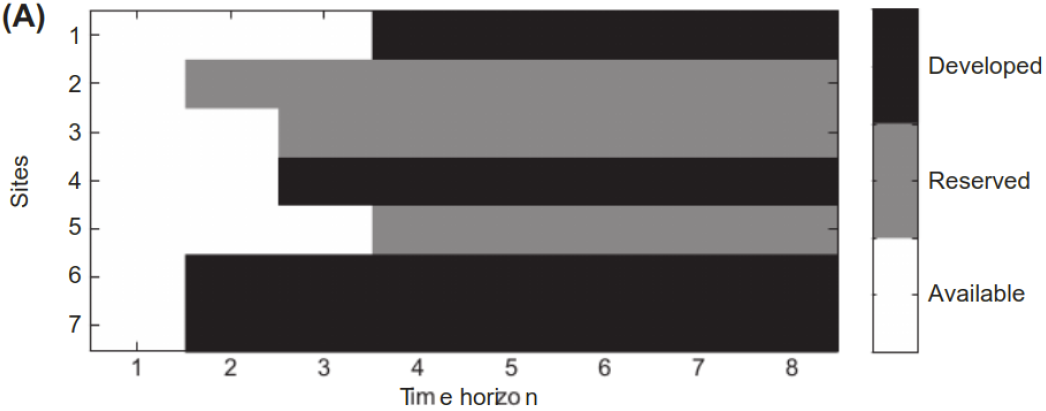
Reserve design problem (Chade`s et al., 2014). The status of 7 sites are represented over a period of 8 time steps. At the beginning (*t* = 1): all sites are available; the optimal action was to reserve site 2; and sites 6 and 7 become developed at the next time step.

## 3 Results

In this section, we report and assess the solutions of the *A*-*K*-MDP algorithm to the previous conservation decision problems. We use MDPtoolbox (Chadès et al., 2014) (available at https://miat.inrae.fr/MDPtoolbox/). The *K*-MDP Matlab package and the conservation problems in .mat format are available as a ZIP file in bit.ly/conservationandai. Supplementary material includes additional information about the performance of the *A*-*K*-MDP algorithm. We set a precision target *p*_*t*_ of 0.0001. Every *K*-MDP is computed in less than a second.

### 3.1 Control of a population: Wolf Culling

For our first example ”Wolf Culling #1” (3000 states, 11 actions), we choose values of *K* ranging from 6 to 3000 states. For *K* = 6 states, i.e., a reduction of 99.83% of the size of the state space, the *A*-*K*-MDP algorithm has a maximum error of only 1%. Figure 7a represents the distribution of the 3000 original states denoting the wolves population, into *K*=6 states. The states are contiguously aggregated according to the number of individuals (See supplementary material for more details). Figure 8 represents the optimal *K*-MDP policy graph, obtained by *A*-*K*-MDP, with 6 states. The *K*-MDP policy graph is consistent with the original optimal policy. However, the original optimal policy would have required 3000 nodes with at most 3000^2^ edges, instead of 6 nodes with at most 2^6^ edges (the reduced optimal policy has exactly 16 edges).

**Figure 7:**
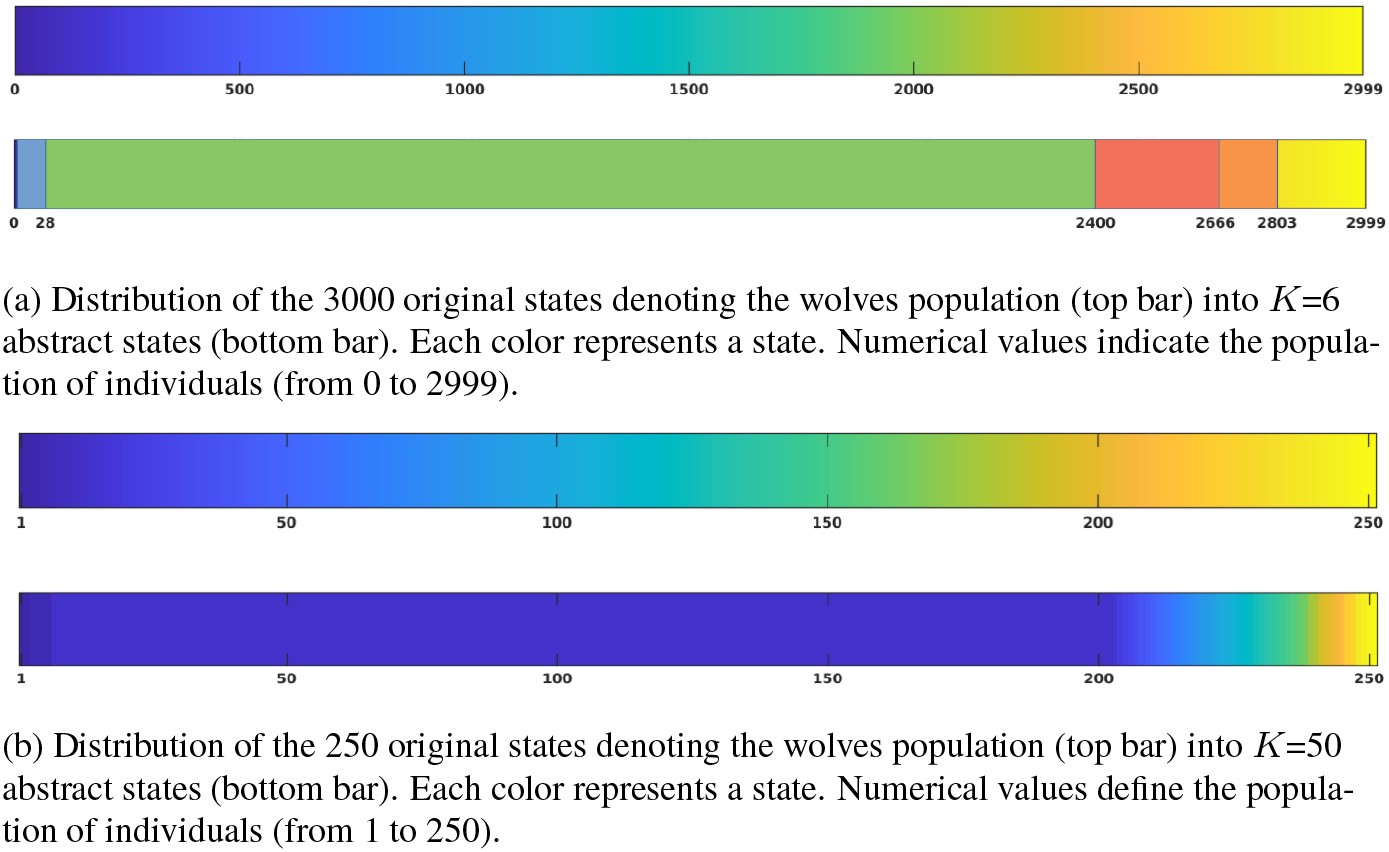
Distribution of original states vs abstract states for two problem instances. *A*-*K*-MDP selects contiguous states to be aggregated.

**Figure 8:**
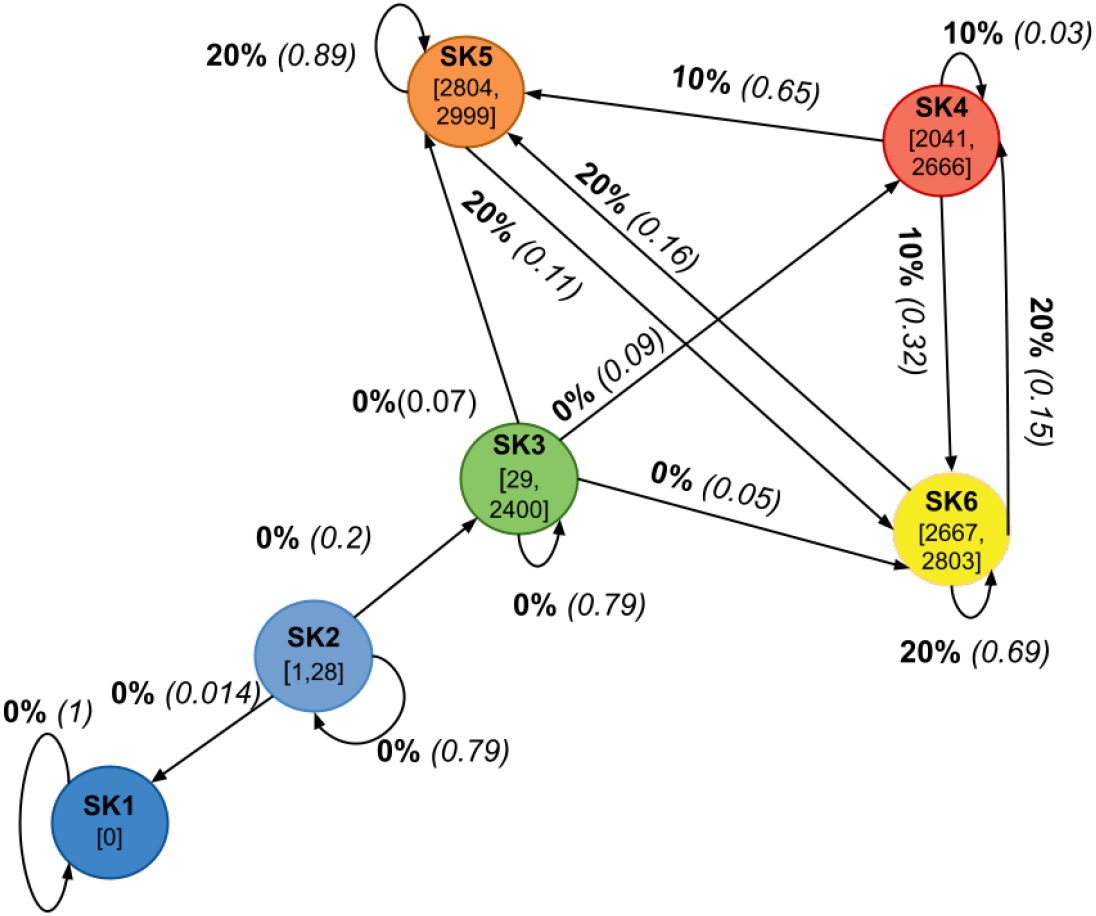
*K*-MDP policy graph for *K* = 6 for the wolf culling problem generated by *A*-*K*-MDP. Nodes represent the abstract states, with 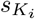 being the abstract state identifier and the interval being the number of individuals represented by that abstract state. Edges are labeled by the optimal harvest rate and its probability to transition to another abstract state.

For our second example, ”Wolf Culling #2” (250 states, 251 actions), we choose values of *K* ranging from 50 to 225 states. For *K* = 50 states, the maximum error was only of 1% in comparison to the original MDP. The *K*-MDP represents a reduction of 80% of the original MDP. The *A*-*K*-MDP algorithm could not find a suitable abstraction with fewer than 50 states. Indeed, the optimal policy uses 50 of the 251 actions, and the 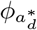 state abstraction requires at least one abstract state per action appearing in the optimal policy. Figure 7b shows the distribution of the 250 original states denoting the wolf population into *K*=50 abstract states. The *A*-*K*-MDP algorithm distributes the 250 original states contiguously into 50 abstract states (see the supplementary material for details on how states have been aggregated).

### 3.2 Recovering two endangered species: Sea otter and northern abalone

For the sea otter and northern abalone problem, we run the *A*-*K*-MDP algorithm for values of *K* ranging from 819 down to 5 states. For *K* = 5 states, we obtain an error of only 1%. This achieves a reduction of 99.39% of the state space. For values of *K* ≤ 4, we could not find a feasible *K*-MDP.

This problem is composed by two state variables, and its aggregated state space is more complex to represent than our previous case studies. Figure 9a represents the density of abalone and abundance of sea otter per each one of the original 819 states. Each square, identified by a color, represents a state. Figure 9b shows the reduced state space representation. There are only 5 different colors representing the *K* = 5 abstract states. A color assigned to a square indicates to what abstract state the original state belongs. For *K* = 5, the state aggregation is:

**Figure 9:**
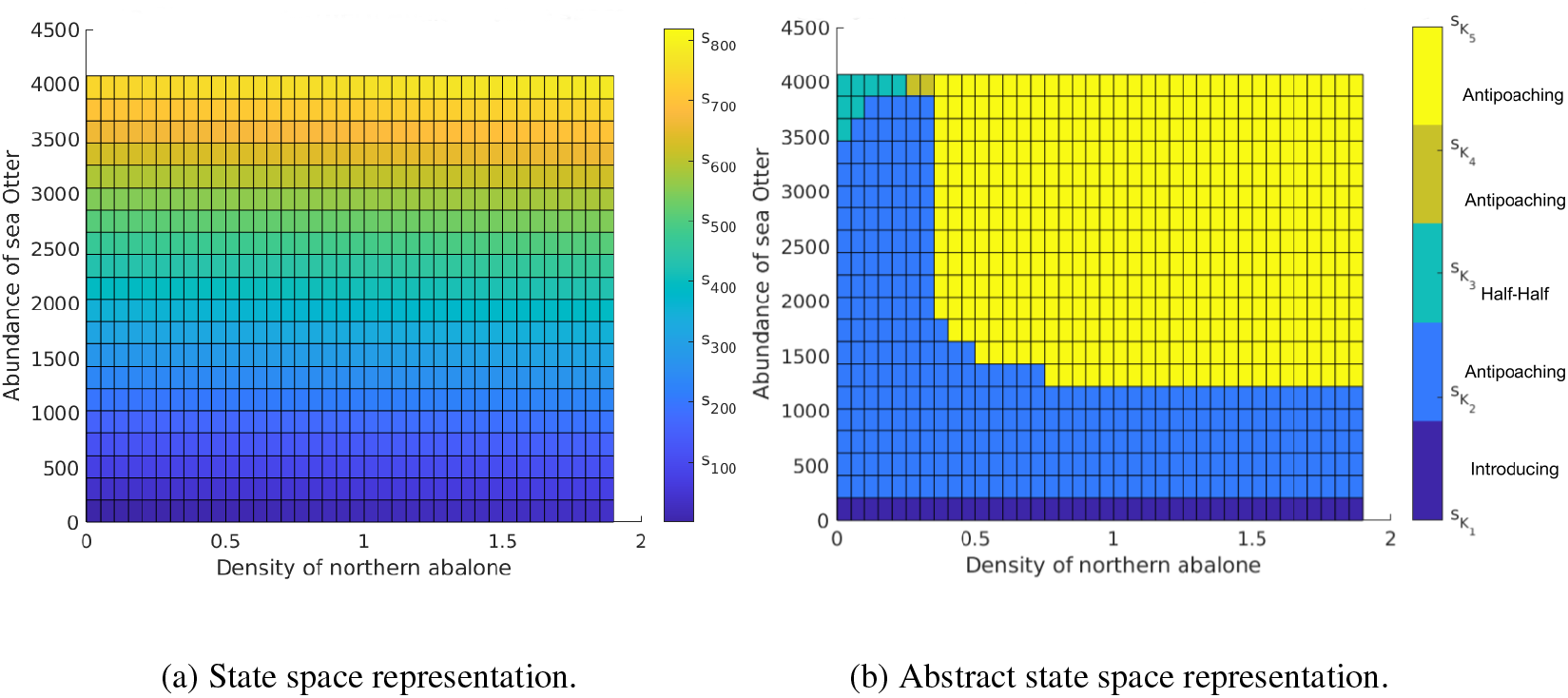
State representation over the abalone density and otter abundance. Each square represents a state with a minimum and maximum abalone density and otter abundance. (a) Each color represents one of the 819 states. (b) Each color represents an abstract state.

- 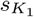 (dark blue): All states with an absence of sea otters. The optimal action is to introduce sea otters.
- 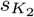 (blue): States where the density of northern abalone is between 0 and 1.9 and the abundance of sea otter is between 250 and 1250, plus states where the density of northern abalone is between 0 and 0.45 and the abundance of sea otter is between 1250 and 3750. The optimal action is antipoaching.
- 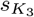 (light blue): States with an abundance of sea otter between 3500 and 4000, and an density of northern abalone between 0.0 and 0.3. The optimal action is half antipoaching and half control sea otters.
- 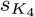 (dark yellow): States where the abundance of sea otter is between 3000 and 4000, and the density of northern abalone is between 0 and 0.3. The optimal action is antipoaching.
- 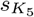 (yellow): States where the abundance of sea otter is between 1250 and 4000, and the density of northern abalone is between 0.35 and 1.9. The optimal action is antipoaching.

Figure 10 represents the optimal *K*-MDP policy graph with only 5 abstract states. The *K*-MDP policy graph is consistent with the original optimal policy. In the absence of sea otters, the optimal action is introducing sea otters; applying antipoaching measures to achieve the short-term recovery target for northern abalone and finally applying half antipoaching and half control sea otters under high predation rates and low density of northern abalone.

**Figure 10:**
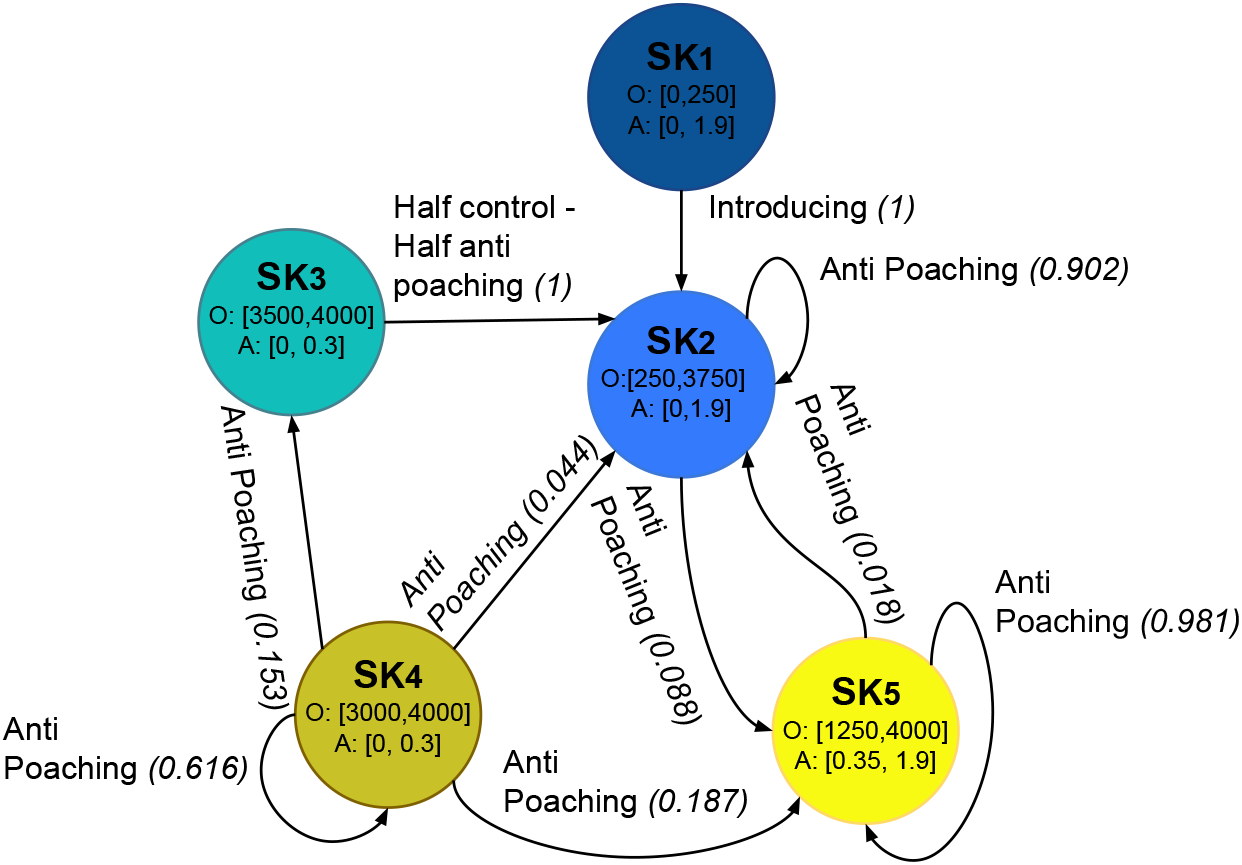
Sea otter and northern abalone optimal *K*-MDP policy graph. Nodes represent the aggregated states, and edges, represent possible outcomes. The starting state is 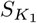. Intervals are approximate.

The original optimal policy would have required 819 nodes with at most 819^2^ edges. The *A*-*K*-MDP algorithm provides a solution with only 5 nodes with at most 5^2^ edges.

### 3.3 Dynamic reserve site selection

For the dynamic reserve site selection problem, we ran the *A*-*K*-MDP algorithm for values of *K* ranging from 729 to 8. The algorithm finds *K*-MDPs with 0% error until *K* = 8. This is a reduction of 98.9% of the state space while retaining 100% of its value. The algorithm cannot find a *K*-MDP when *K <* 8. Given the multivariate nature of this problem, with 6 variables denoting each one of the sites, finding a suitable and interpretable representation of the resulting *K* = 8 *K*-MDP and the resulting policy graph required additional steps.

Analysing the grouping of original states into abstract states, we identified 3 patterns within the abstract states:

- Abstract state 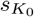 represents the states where all sites are available.
- Abstract state 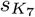 represents the states where all sites are developed.
- Interestingly, abstract states 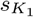 to 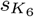 group states together that have a unique site that has not been developed. For example, 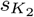 does not contain states where the value of the state variable denoting site 1 is developed. Formally, 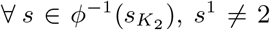, where *s*^1^ denotes the state variable for site 1. While this is an interesting finding, this rule does not allow us to uniquely identify which original states belong to which abstract states. For example, a small proportion of states with site 1 taking value available or reserved are not in 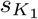.

Figure 11 represents the optimal *K*-MDP policy with only 8 states. For convenience, we only show edges for transition probabilities greater than 0.1. The original policy graph would have required 729 nodes and at most 729^2^ edges. Using *A*-*K*-MDP, we can compute a policy graph with only 8 nodes and 19 edges. Evaluating the policy graph, the optimal action to apply in each abstract state is to reserve the site that is not currently developed in that abstract state, except for abstract states 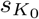 (Reserve site 3) and 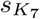 (Reserve site 1).

**Figure 11:**
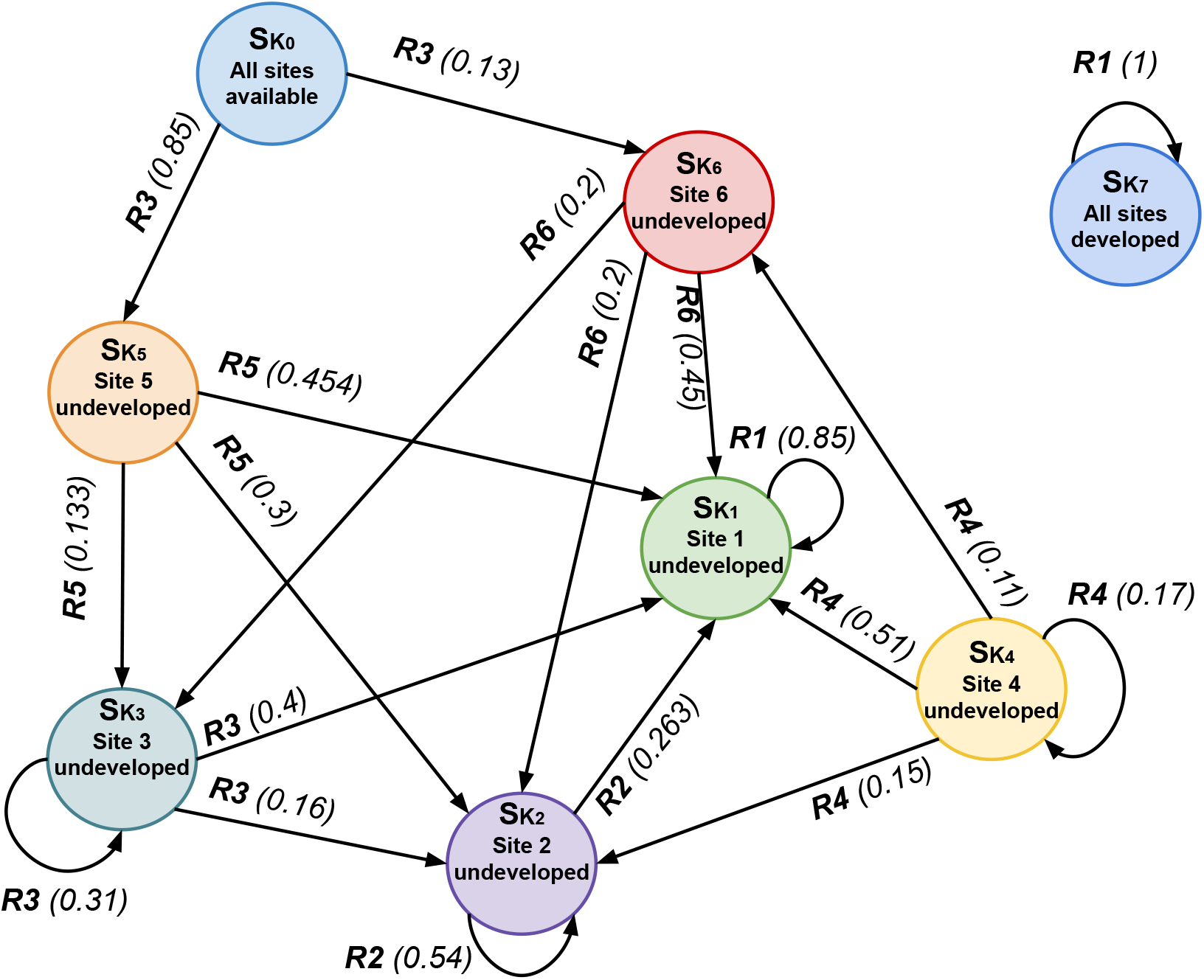
*K*-MDP policy graph for *K* = 8 for the dynamic reserve site problem. *R*_*i*_(*p*) denotes the site to reserve and the transition probability. For simplicity, we removed edges with transition probability less than 0.1. Edges to abstract state 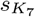 have been deleted because transitions to that state had probabilities lower than 0.1, which means that 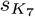”All sites developed” is an unlikely state to occur in the system. The action R1 in 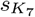 is an arbitrary choice and has no effect.

To better understand the state aggregation and the optimal reduced policy, recall that the objective of this problem is to maximise the number of unique species protected, and a species-by-site matrix generated for this problem is given in Table 4. The sites-by-species matrix shows the distribution of the 7 species along the 6 sites. The optimal action to apply in abstract state 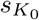, when all sites are available, is to reserve site 3. This is because reserving site 3 results in five of the seven species being protected at once (species 1, 4, 5, 6, and 7). The resulting state is either 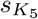 (site 5 non developed) or 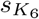 (site 6 non developed), with their optimal actions being reserving site 5 and reserving site 6, respectively. Reserving any of these two sites will protect the remaining unprotected species (species 2 and 3), achieving the objective of protecting the total number of species (7).

**Table 4:**
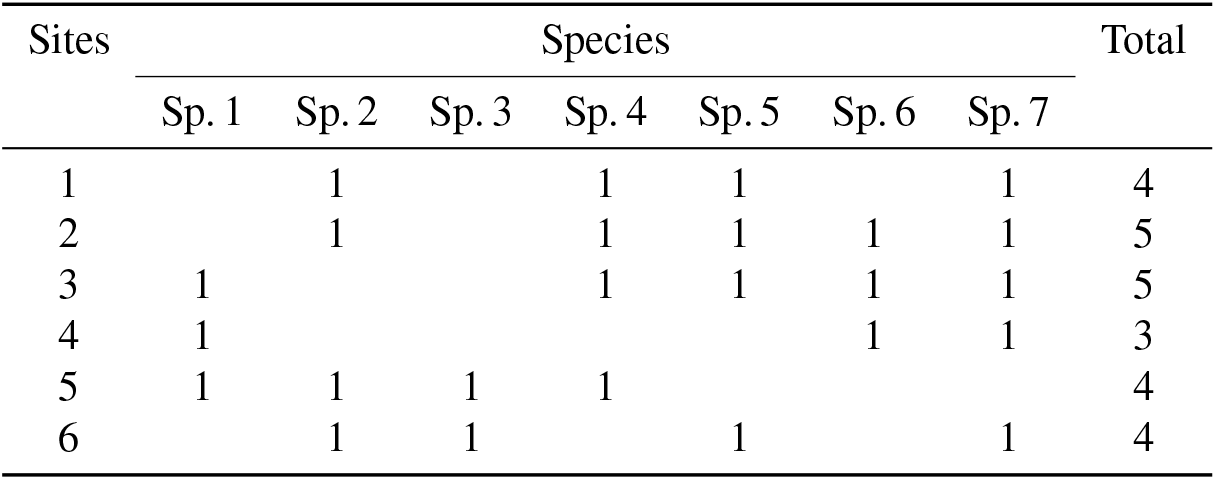
Sites-by-species matrix for the dynamic reserve site selection case study. Rows represent the 6 sites and columns 1 to 7 represent the 7 species to be protected. Last column shows the total number of species per site.

| Sites | Species |  |  |  |  |  |  | Total |
| --- | --- | --- | --- | --- | --- | --- | --- | --- |
|  | Sp. 1 | Sp. 2 | Sp. 3 | Sp. 4 | Sp. 5 | Sp. 6 | Sp. 7 |  |
| 1 |  | 1 |  | 1 | 1 |  | 1 | 4 |
| 2 |  | 1 |  | 1 | 1 | 1 | 1 | 5 |
| 3 | 1 |  |  | 1 | 1 | 1 | 1 | 5 |
| 4 | 1 |  |  |  |  | 1 | 1 | 3 |
| 5 | 1 | 1 | 1 | 1 |  |  |  | 4 |
| 6 |  | 1 | 1 |  | 1 |  | 1 | 4 |

We are not able to provide a visual representation of the original optimal policy graph given its size of 729 nodes and up to 729^2^ edges, but, thanks to the *A*-*K*-MDP algorithm, we have been able to provide and interpret a reduced policy graph.

Optimal actions and transitions between other abstract states of the policy graph can similarly be explained using the information given by the sites-by-species matrix.

## 4 Discussion

MDPs are increasingly applied to model ecological and natural resource management decision-making problems. We propose an approach to reduce the size of MDPs, and thus of the resulting policies, to facilitate the extraction of meaningful recommendations. We have shown that the *A*-*K*-MDP algorithm can compute an MDP with a small number of states (*K*-MDP) from an original MDP. Given an MDP model and a constraint on the number of states *K*, the *A*-*K*-MDP algorithm computes a simpler MDP with at most *K* states, while minimizing the difference between the original optimal MDP value function and the reduced optimal *K*-MDP value function.

We assessed the *K*-MDP approach on three case studies of increasing complexity: from one state variable (Marescot et al., 2013) to six state variables (Sabbadin et al., 2007). For reproducibility, we provide the code as a Matlab package and the MDP models of our case studies. For each of these illustrative problems, we successfully reduced their state spaces to just a small number (*K*) of meaningful abstract states at a small performance cost. Our experiments show that the *A*-*K*-MDP algorithm achieves a considerable reduction in the number of states while producing policies that at least retain 98% of the original value. By reducing the size of the resulting solution, it was always easier to represent and interpret the solutions.

While our algorithm achieves remarkable reduction in the size of the state space with little performance loss, it is worth noting that not all problems would require a reduction of their state space. For example, our first case study, tackling the control of a population, has only one state variable. The optimal policy can be represented by a simple graph (Figure 4) and it is already easy to interpret. Nonetheless, our approach does provide a very compact representation of the policy graph (Figure 8). In problems with two state variables, such as the second case study, the *A*-*K*-MDP algorithm clearly facilitates the interpretation of the optimal solution. For problems with more state variables, such as the dynamic reserve site selection problem, while our approach tremendously simplified the original problem, we had to perform additional analysis to determine a unique property (i.e., site non-developed) to identify the meaning of the computed abstract states. Future research should aim to further facilitate the identification of explainable features of the *K*-MDP approach.

The *A*-*K*-MDP algorithm requires first solving an MDP to be able to apply the abstraction function (equation 5), because the aim was not to solve MDPs but rather to provide a more compact representation of the solution. Other aggregation criteria can be applied, such as qualitative constraints based on the problem at hand. Ferrer-Mestres and Chadès (2021) explored selecting aggregation rules designed directly by the end-user rather than the presented state abstraction functions. It creates more meaningful abstractions from the human-operator’s point of view at a small loss of performance.

Broadening the scope of this work, MDPs can also be thought of as a quantitative framework for state and transition models (STMs) (Bestelmeyer et al., 2017). Indeed, the state definition and their transitions given a management action fit the definition of STMs. In this manuscript, we have not explored how the *K*-MDP approaches could also help increase the interpretability of STM models. This is because although STMs are used to guide decision-making, they do not specify reward functions that are required to define MDPs and optimise the decision-making processes. Future research could explore how state abstraction functions could be applied to aggregate states without specification of reward elements.

Modeling a problem as an MDP is an art. When designing an MDP, an important modeling decision is to determine the number of state variables and their discrete values. The *A*-*K*-MDP algorithm allows the designer to instead start with a large state space with many state variables and many values and let the algorithm determine how best to abstract that state space while retaining the distinctions that are important for achieving good policy performance. We believe that the *A*-*K*-MDP algorithm has the potential to become a new standard tool to increase interpretability of AI decision tools and increase adoption for better on-ground ecological management outcomes.

## 6 Data Availability

All code and data relevant to this study is available as a ZIP file and can be found in bit.ly/conservationandai.

## References

Abel, D., Arumugam, D., Lehnert, L. and Littman, M. (2018) State abstractions for lifelong reinforcement learning. In Proceedings of the 35th International Conference on Machine Learning (eds. J. Dy and A. Krause), vol. 80 of Proceedings of Machine Learning Research, 10–19. PMLR. URL: https://proceedings.mlr.press/v80/abel18a.html.

Abel, D., Hershkowitz, D. and Littman, M. (2016) Near optimal behavior via approximate state abstraction. In Proceedings of The 33rd International Conference on Machine Learning (eds. M. F. Balcan and K. Q. Weinberger), vol. 48 of Proceedings of Machine Learning Research, 2915–2923. New York, New York, USA: PMLR. URL: https://proceedings.mlr.press/v48/abel16.html.

Bellman, R. (1957) Dynamic programming. Princeton University Press, John Wiley & Sons.

Bestelmeyer, B. T., Ash, A., Brown, J. R., Densambuu, B., Fernández-Giménez, M., Johanson, J., Levi, M., Lopez, D., Peinetti, R., Rumpff, L. et al. (2017) State and transition models: theory, applications, and challenges. In Rangeland systems, 303–345. Springer, Cham.

Boettiger, C., Mangel, M. and Munch, S. (2015) Avoiding tipping points in fisheries management through Gaussian process dynamic programming. Proceedings of the Royal Society B: Biological Sciences, 282, 20141631.

Brechtel, S., Gindele, T. and Dillmann, R. (2011) Probabilistic MDP-behavior planning for cars. In Proceedings of the 14th International IEEE Conference on Intelligent Transportation Systems (ITSC), 1537–1542. IEEE.

Chadès, I., Chapron, G., Cros, M.-J., Garcia, F. and Sabbadin, R. (2014) MDPtoolbox: a multiplatform toolbox to solve stochastic dynamic programming problems. Ecography, 37, 916–920.

Chades, I., Curtis, J. M. R. and Martin, T. G. (2012) Setting realistic recovery targets for two interacting endangered species, sea otter and northern abalone. Conservation Biology, 26, 1016–1025. URL: http://www.jstor.org/stable/23360117.

Collazo, J. A., Fackler, P. L., Pacifici, K., White Jr, T. H., Llerandi-Roman, I. and Dinsmore, S. J. (2013) Optimal allocation of captive-reared Puerto Rican parrots: Decisions when divergent dynamics characterize managed populations. The Journal of wildlife management, 77, 1124–1134.

Costello, C. and Polasky, S. (2004) Dynamic reserve site selection. Resource and Energy Economics, 26, 157–174.

Dean, T. and Givan, R. (1997) Model minimization in Markov decision processes. In Proceedings of the Fourteenth National Conference on Artificial Intelligence and Ninth Conference on Innovative Applications of Artificial Intelligence (AAAI/IAAI), 106–111.

Dean, T., Givan, R. and Leach, S. (1997) Model reduction techniques for computing approximately optimal solutions for Markov decision processes. In Proceedings of the Thirteenth Conference on Uncertainty in Artificial Intelligence (UAI), 124–131. Morgan Kaufmann Publishers Inc.

Fackler, P. (2011) MDPSolve. URL: https://github.com/PaulFackler/MDPSolve.

Ferrer-Mestres, J. and Chadès, I. (2021) Solving constrained K-Markov decision processes. In Proceedings of the 24th International Congress on Modelling and Simulation (MODSIM).

Ferrer-Mestres, J., Dietterich, T. G., Buffet, O. and Chadès, I. (2020) Solving K-MDPs. In Proceedings of the International Conference on Automated Planning and Scheduling (ICAPS), vol. 30, 110–118.

Johnson, F. A., Breininger, D. R., Duncan, B. W., Nichols, J. D., Runge, M. C. and Williams, B. K. (2011) A Markov decision process for managing habitat for Florida scrub-jays. Journal of Fish and Wildlife Management, 2, 234–246.

Kober, J., Bagnell, J. A. and Peters, J. (2013) Reinforcement learning in robotics: A survey. The International Journal of Robotics Research, 32, 1238–1274.

Koopmans, T. C. (1960) Stationary ordinal utility and impatience. Econometrica: Journal of the Econometric Society, 287–309.

Li, L., Walsh, T. J. and Littman, M. L. (2006) Towards a unified theory of state abstraction for MDPs. In International Symposium on Artificial Intelligence and Mathematics (ISAIM).

Mangel, M., Clark, C. W. et al. (1988) Dynamic modeling in behavioral ecology, vol. 63. Princeton University Press.

Marescot, L., Chapron, G., Chades, I., Fackler, P. L., Duchamp, C., Marboutin, E. and Gimenez, O. (2013) Complex decisions made simple: a primer on stochastic dynamic programming. Methods in Ecology and Evolution, 4, 872–884.

McCarthy, M. A., Possingham, H. P. and Gill, A. M. (2001) Using stochastic dynamic programming to determine optimal fire management for Banksia ornata. Journal of Applied Ecology, 38, 585–592.

Memarzadeh, M., Britten, G. L., Worm, B. and Boettiger, C. (2019) Rebuilding global fisheries under uncertainty. Proceedings of the National Academy of Sciences, 116, 15985–15990.

Milner-Gulland, E. (1997) A stochastic dynamic programming model for the management of the saiga antelope. Ecological Applications, 7, 130–142.

Moore, J. L., Rout, T. M., Hauser, C. E., Moro, D., Jones, M., Wilcox, C. and Possingham, H. P. (2010) Protecting islands from pest invasion: optimal allocation of biosecurity resources between quarantine and surveillance. Biological Conservation, 143, 1068–1078.

Nicol, S. and Chadès, I. (2012) Which states matter? An application of an intelligent discretization method to solve a continuous POMDP in conservation biology. PLoS ONE, 7. URL: 10.1371/journal.pone.0028993.

Péron, M., Jansen, C. C., Mantyka-Pringle, C., Nicol, S., Schellhorn, N. A., Becker, K. H. and Chadès, I. (2017) Selecting simultaneous actions of different durations to optimally manage an ecological network. Methods in Ecology and Evolution, 8, 1332–1341.

Petrik, M. and Luss, R. (2016) Interpretable policies for dynamic product recommendations. In Proceedings of the Thirty-Second Conference on Uncertainty in Artificial Intelligence (UAI).

Pozzi, M., Memarzadeh, M. and Klima, K. (2017) Hidden-model processes for adaptive management under uncertain climate change. Journal of Infrastructure Systems, 23, 04017022.

Puterman, M. L. (2014) Markov decision processes: discrete stochastic dynamic programming. John Wiley & Sons.

Reeson, A. and Dunstall, S. (2009) Behavioural economics and complex decision-making. Victoria: CSIRO.

Sabbadin, R., Spring, D. and Rabier, C.-E. (2007) Dynamic reserve site selection under contagion risk of deforestation. Ecological Modelling, 201, 75–81.

Schwarz, L. K., McHuron, E., Mangel, M., Wells, R. S. and Costa, D. P. (2016) Stochastic dynamic programming: An approach for modelling the population consequences of disturbance due to lost foraging opportunities. In Proceedings of Meetings on Acoustics 4ENAL, vol. 27, 040004. Acoustical Society of America.

Sigaud, O. and Buffet, O. (2013) Markov decision processes in artificial intelligence. John Wiley & Sons.

Tulloch, V. J., Tulloch, A. I., Visconti, P., Halpern, B. S., Watson, J. E., Evans, M. C., Auerbach, N. A., Barnes, M., Beger, M., Chadès, I. et al. (2015) Why do we map threats? Linking threat mapping with actions to make better conservation decisions. Frontiers in Ecology and the Environment, 13, 91–99.

Venner, S., Chadès, I., Bel-Venner, M.-C., Pasquet, A., Charpillet, F. and Leborgne, R. (2006) Dynamic optimization over infinite-time horizon: web-building strategy in an orb-weaving spider as a case study. Journal of theoretical biology, 241, 725–733.

Westphal, M. I., Pickett, M., Getz, W. M. and Possingham, H. P. (2003) The use of stochastic dynamic programming in optimal landscape reconstruction for metapopulations. Ecological Applications, 13, 543–555.

Wilson, K. A., McBride, M. F., Bode, M. and Possingham, H. P. (2006) Prioritizing global conservation efforts. Nature, 440, 337–340.

