## Supplementary Information for "Interpretable Solutions for Stochastic Dynamic Programming"

### 1 Supplementary material

This supplementary material provides an extended evaluation of the empirical results, additional details of state abstraction functions and the algorithm for building $K$-MDPs.

#### 1.1 Detailed Experimental Results

We ran our experiments on an Intel i7-8650U CPU, 1.90 GHz, with 16 GB RAM and Ubuntu 18.04. Experiments were conducted using the *MDPToolbox* (Chadès et al., 2014), MATLAB (R2019b) and the $K$-MDP Matlab package (It can be found here). We acknowledge there are other available MDP solvers (Fackler, 2011; Kurniawati et al., 2008). Each problem is modeled as an MDP. All problems are accessible via the attached ZIP file. The problems are comprised of two files: i) A transition matrix $P$ of size $|S| \times |S| \times |A|$, denoting the dynamics of the system, where $|S|$ is the number of states and $|A|$ is the number of applicable actions; and ii) A reward matrix $R$ of size $|S| \times |A|$ or size $|S| \times |S| \times |A|$. In this section, we provide an extended description of the abstract states for the wolf culling and the dynamic site selection case studies. We assume that the weights in transitions and rewards follow a uniform probability distribution across the original states: $\forall s \in \phi^{-1}(s_K), \omega(s) = \frac{1}{|\phi^{-1}(s_K)|}$.

##### 1.1.1 Control of a population: Wolf culling problem

This problem was published by Marescot et al. (2013).

For $K = 6$, the abstract states contiguously aggregate the ground states according to the population of individuals:

- Abstract state $s_{K_1}$ aggregates state $s_1$: 0 individuals. The optimal action is 0% culling;
- Abstract $s_{K_2}$ aggregates states from $s_2$ to $s_{29}$: From 1 to 28 individuals. The optimal action is 0% culling;
- Abstract state $s_{K_3}$ aggregates states from $s_{30}$ to $s_{2401}$: From 29 to 2400 individuals. The optimal action is 0% culling;
- Abstract state $s_{K_4}$ aggregates states from $s_{2042}$ to $s_{2667}$: From 2401 to to 2666. The optimal action is 10% culling;

- Abstract state $s_{K_5}$ aggregates states from $s_{2805}$ to $s_{3000}$: From 2804 to 2999 individuals. The optimal action is 20% culling;
- Abstract state $s_{K_6}$ aggregates states from $s_{2668}$ to $s_{2804}$: From 2667 to 2803 individuals. The optimal action is 20% culling.

For $K$=6, the $A$-$K$-MDP algorithm has an error of only $1\%$. This is a reduction of a 99.83% of the state space.

For $K = 50$, the state aggregation and optimal policy is:

- Abstract state $s_{K_1}$ aggregates state $s_1$: 0 individuals. The optimal policy is 0% culling.
- Abstract state $s_{K_2}$ aggregates states from $s_2$ to $s_5$: From 1 to 4 individuals. The optimal policy is 0% culling.
- Abstract state $s_{K_3}$ aggregates states from $s_6$ to $s_{201}$: From 5 to 200 individuals. The optimal policy is 0% culling.
- Abstract states from $s_{K_4}$ to $s_{K_{39}}$ aggregate states from $s_{202}$ to $s_{237}$ respectively: From 201 to 236 individuals. The optimal policy is from 2% to 39% culling respectively.
- Abstract state $s_{K_{40}}$ aggregates states $s_{238}$ and $s_{239}$: 237 and 238 individuals. The optimal policy is 40% culling.
- Abstract states from $s_{K_{41}}$ to $s_{K_{45}}$ aggregate states from $s_{240}$ to $s_{244}$ respectively: From 239 to 243 individuals. The optimal policy is from 41% to 45% culling respectively.
- Abstract state $s_{K_46}$ aggregates states $s_{245}$ and $s_{246}$: 244 and 245 individuals. The optimal policy is 46% culling.
- Abstract states from $s_{K_{47}}$ to $s_{K_{50}}$ aggregate states from $s_{247}$ to $s_{250}$ respectively: From 246 to 249 individuals. The optimal policy is from 47% to 50% culling respectively.

For $K$=50, the $A$-$K$-MDP algorithm has an error of only $1\%$. This $K$-MDP represents a reduction of 80% of the state space. Our algorithm is not able to find $K$-MDPs for $K \leq 50$. This is because $A$-$K$-MDP can only find state abstractions of size $K$ equal or greater to the number of optimal actions in the optimal policy $\pi^*$. We acknowledge that other state abstractions functions and $K$-MDP algorithms could find feasible abstractions with $K \leq 50$ at a cost on the loss of performance.

##### 1.1.2 Dynamic reserve site selection

The dynamic reserve site selection problem poses the greatest challenge in terms of visual representation and interpretation. This problem has six state variables representing the state of each one of the seven sites (available, reserved or developed). The $A$-$K$-MDP algorithm performs similarly to those problems with only one state variable, with an error close to 0% for most values of $K$. For $K = 8$, the resulting state aggregation is:

- Abstract state $s_{K_0}$ aggregates the unique state where all sites are available. The optimal action is reserving site 3. By reserving site 3, 5 different species are protected (species 1, 4, 5, 6, and 7).
- Abstract state $s_{K_1}$ aggregates states where site 1 is not developed. The optimal action is reserving site 1.
- Abstract state $s_{K_2}$ aggregates states where site 2 is not developed. The optimal action is reserving site 2.
- Abstract state $s_{K_3}$ aggregates states where site 3 is not developed. The optimal action is reserving site 3.
- Abstract state $s_{K_4}$ aggregates states where site 4 is not developed. The optimal action is reserving site 4.
- Abstract state $s_{K_5}$ aggregates states where site 5 is not developed. The optimal action is reserving site 5.
- Abstract state $s_{K_6}$ aggregates states where site 6 is not developed. The optimal action is reserving site 6.
- Abstract state $s_{K_7}$ aggregates the unique state where all sites are developed. The optimal action is irrelevant. The optimal value at this state is always 0.

#### 1.2 State abstraction functions

The original motivation for introducing state abstractions was to speed up learning and optimization (Dean and Givan, 1997; Li et al., 2006). In this paper, our goal is to improve the interpretability of MDP models and solutions. We define the predicate $p(s_1, s_2)$ to be true when $s_1$

and $s_2$ can be abstracted. Then, a state abstraction function $\phi$ satisfies:

$$(\phi(s_1) = \phi(s_2)) \implies p(s_1, s_2). \tag{1}$$

State abstraction functions have some desirable properties and can be:

- **Exact:** There is no difference between two states given a metric. The loss of performance is 0.
- **Approximate:** There is a difference between two states and the loss of performance is bounded by a parameter $\epsilon$.
- **Non-transitive:** Two possible state abstractions between $s_1$ and $s_2$ ($p(s_1, s_2)$) and between $s_2$ and $s_3$ ($p(s_2, s_3)$) do not imply that there exists an abstraction between $s_1, s_3$ ($p(s_1, s_3)$).
- **Transitive:** Two state abstractions $p(s_1, s_2)$ and $p(s_2, s_3)$ together imply $p(s_1, s_3)$.

We focus on approximate transitive state abstraction functions because of the advantages that they offer in front of exact non-transitive abstractions. First, approximate state abstractions allow greater degrees of aggregation between states while exact state abstraction functions aggregate states that are exactly equal given a metric (Li et al., 2006; Dean and Givan, 1997), but this very strong requirement limits our ability to aggregate states. Second, with transitive state abstraction predicates we can reduce many calculations.

#### 1.3 Algorithms to solve $K$-MDPs

This section includes the pseudo-code for the *BUILD-K-MDP* procedure (Algorithm 1). This function takes as input an MDP $M$, an abstract state space $S_K$ produced by our algorithm and the abstraction function $A$-$K$-MDP. Then the procedure computes the weights of all states $s \in \phi^{-1}(s_K)$ (line 3), the corresponding reward function $r_K$ (line 5) and transition function $T_K$ (line 9). For simplicity, we assumed that, for any given abstract fully observable state $s_K$, if the number of original fully observable states aggregated to $s_K$ is $|\phi^{-1}(s_K)|$, then the weight of each $s \in \phi^{-1}(s_K)$ is uniformly distributed: $\omega(s) = \frac{1}{|\phi^{-1}(s_K)|}$.

**Algorithm 1** *BUILD-$K$-MDP*

**Require:** $M = \langle S, A, T, r, H, \gamma \rangle, S_K, \phi$

**for** $s_K \in S_K$ **do**<br>
  **for** $s \in \phi^{-1}(s_K)$ **do**<br>
    $\omega(s) \leftarrow$ *computeWeights*$(\phi, s_K)$<br>
  **end for**<br>
  $r_K(s_K, a) \leftarrow \sum_{s \in \phi^{-1}(s_K)} r(s, a)\omega(s)$<br>
  **for** $s'_K \in S_K$ **do**<br>
    **for** $a \in A$ **do**<br>
      $T_K(s_K, a, s'_K) \leftarrow$<br>
          $\sum_{s \in \phi^{-1}(s_K)} \sum_{s' \in \phi^{-1}(s'_K)} T(s, a, s')\omega(s)$<br>
    **end for**<br>
  **end for**<br>
**end for**<br>
$M_K = \langle S_K, A, T_K, r_K, H, \gamma \rangle$<br>
**return** $M_K$

### 2 Notation

Table 1 shows the different notations used in MDPs and $K$-MDPs.

Table 1: Notations used in MDP and $K$-MDP models.

| Variable | MDP | $K$-MDP |
|---|---|---|
| State space | $S$ | $S_K$ |
| Action space | $A$ | $A$ |
| Transition | $T(s', a, s)$ | $T_K(s'_K, a, s_K)$ |
| Reward | $r(s, a)$ | $r_K(s_K, a)$ |
| Optimal policy | $\pi^*$ | $\pi^*_K$ |
| Optimal value | $V^*$ | $V_\phi^{\pi^*_K}$ |

### References


Chadès, I., Chapron, G., Cros, M.-J., Garcia, F. and Sabbadin, R. (2014) MDPtoolbox: a multi-platform toolbox to solve stochastic dynamic programming problems. *Ecography*, **37**, 916–920.

Dean, T. and Givan, R. (1997) Model minimization in Markov decision processes. In *Proceed-*

*ings of the Fourteenth National Conference on Artificial Intelligence and Ninth Innovative Applications of Artificial Intelligence Conference (AAAI/IAAI)*, 106–111.

Fackler, P. (2011) MDPSolve. URL: `https://github.com/PaulFackler/MDPSolve`.

Kurniawati, H., Hsu, D. and Lee, W. S. (2008) SARSOP: Efficient point-based POMDP planning by approximating optimally reachable belief spaces. In *In Proc. Robotics: Science and Systems*.

Li, L., Walsh, T. J. and Littman, M. L. (2006) Towards a unified theory of state abstraction for MDPs. In *Proceedings of the International Symposium on Artificial Intelligence and Mathematics (ISAIM)*.

Marescot, L., Chapron, G., Chades, I., Fackler, P. L., Duchamp, C., Marboutin, E. and Gimenez, O. (2013) Complex decisions made simple: a primer on stochastic dynamic programming. *Methods in Ecology and Evolution*, **4**, 872–884.
